# Measuring the co-localization and dynamics of mobile proteins in live cells undergoing signaling responses

**DOI:** 10.1101/2022.10.17.511423

**Authors:** Sarah A. Shelby, Thomas R. Shaw, Sarah L. Veatch

## Abstract

Single molecule imaging in live cells enables the study of protein interactions and dynamics as they participate in signaling processes. When combined with fluorophores that stochastically transition between fluorescent and reversible dark states, as in super-resolution localization imaging, labeled molecules can be visualized in single cells over time. This improvement in sampling enables the study of extended cellular responses at the resolution of single molecule localization. This chapter provides optimized experimental and analytical methods used to quantify protein interactions and dynamics within the membranes of adhered live cells. Importantly, the use of pair-correlation functions resolved in both space and time allows researchers to probe interactions between proteins on biologically relevant distance and time-scales, even though fluorescence localization methods typically require long times to assemble well-sampled reconstructed images. We describe an application of this approach to measure protein interactions in B cell receptor signaling and include sample analysis code for post-processing of imaging data. These methods are quantitative, sensitive, and broadly applicable to a range of signaling systems.

## 1. Introduction

The invention of super-resolution microscopy has revolutionized biological imaging, allowing researchers to directly visualize the organization and interactions of proteins on distance-scales relevant to cellular processes. In the field of immune receptor signaling, super-resolution imaging methods have, for example, helped to define cellular contacts between immune cells and antigen presenting cells as well as the composition of signaling platforms formed after immune receptors engage with antigens or cross-linkers [1–7]. Single molecule fluorescence localization microscopy (SMLM) is one method within the broader class of super-resolution imaging that achieves high spatial resolution by imaging sparsely distributed fluorophores over time [8–11]. Fluorophores labeling proteins of interest stochastically transition between emitting and transiently dark states, allowing for the ensemble of molecules to be fully sampled over time. Single and multi-color SMLM has been used to image components of the immune response in a wide variety of contexts [12–21]. Our own work has largely focused on probing early events in B cell receptor signaling [22–24].

An important limitation of SMLM, especially when applied to living systems, is that it typically takes seconds to minutes to acquire an image because many (typically 100s to 1000s) individual images of single molecules are needed to reconstruct a well sampled super-resolved image. Over these extended time-periods, molecules of interest tend to diffuse over distances much larger than the biological structures under investigation, washing out the super-resolved information researchers seek to extract. One approach to overcoming this limitation is to focus on systems with slow dynamics [25, 26], or to push the limits of camera frame-rates and fluorophore photo-physics [27]. In our own past work, we have sought to overcome this limitation by extracting information from SMLM datasets without reconstructing images, by interrogating the relative distributions of molecules with pair-correlation functions.

Pair correlation functions are distributions of pair-wise distances between localized molecules imaged over time. Correlation functions can take the form of the autocorrelation, which tabulates pairwise distances between molecules of the same type, or the cross-correlations, which tabulates pairwise distances between molecules of the different types. Autocorrelation functions provide information on how labeled proteins move and how they self-associate, while cross-correlation functions probe the colocalization of proteins, as well as the dynamics and effective energy of these associations [15, 28–33]. Pair-correlation methods can also be used as part of a general image processing pipeline to enhance or characterize SMLM datasets [34, 35].

Previously we have described how a steady-state cross-correlation can extract information relevant to interactions between proteins in live cells undergoing signaling responses after the B cell receptor is engaged with a multivalent cross-linker [22, 31]. Since then, we have further optimized and extended these approaches, enabling the detection of weak signals over time [24]. This protocol chapter describes optimized experimental approaches including sample preparation, fluorescent probe selection, and microscopy imaging methods. We also include protocols for single fluorophore localization in raw images, the evaluation of properly normalized pair correlation functions, and use of correlation functions to characterize colocalization and molecular motions in live cells. Analysis code is provided to facilitate the application of these methods by other groups.

## 2. Materials

### 2.1 Cell culture and transfection

1. Cells and culture medium. The imaging methods described here can be used to study interactions and dynamics of plasma membrane signaling proteins in a variety of cell types. The only requirements are a relatively flat cell membrane surface for imaging through cell adhesion to a bare glass or functionalized coverslip, and that proteins of interest can be labeled with SMLM-compatible fluorescent tags. For our studies of the B cell receptor, we use CH27 mouse B cells [36]. CH27 cells are maintained in culture in growth media at 37°C and 5% CO2 as previously reported [31].
2. Reagents and/or equipment for transient transfection of plasmid DNA. Any system for transient transfection can be used that allows for robust expression of fluorescent protein constructs of interest in the cell line under study. We use a Lonza 4D Nucleofector X unit following the manufacturer’s protocols for electroporation of CH27 B cells.
3. Fluorescent fusion constructs: proteins of interest can be labeled in imaging experiments through expression of proteins fused to a super-resolution compatible fluorescent protein. We typically use the photoswitchable protein mEos3.2. for our experiments [37]. Generally, DNA encoding proteins of interest are cloned into mEos3.2-C1 or mEos3.2-N1 vectors under a CMV promoter.

### 2.2 Dye conjugation of labeling proteins

Reactive dyes and unlabeled monomeric labeling reagents: In-house dye conjugation of labeling proteins (e.g. antibody Fab fragments) is often required. For situations where protein clustering is to be avoided or controlled, labeling protein-target binding should be monovalent.

1. Unconjugated purified proteins that bind cell surface proteins of interest. We label IgM BCR using goat anti-Mouse IgM(µ) Fab fragments (Jackson ImmunoResearch 115-007-020).
2. Amine-reactive dye succinimidyl esters. Solubilize dye succinimidyl esters in anhydrous DMSO at 10mM. Store stock solution aliquots at −80°C with desiccation, avoid freeze thaw. We use biotin and silicon rhodamine (SiR) NHS esters (Invitrogen: B1582, Spirochrome: SC003) to label and functionalize anti-BCR Fab fragments.
3. Conjugation reaction buffer: 0.01M NaH2PO4 with 0.01M NaH2CO3 at pH 8.5.
4. Gel filtration column e.g. GE Healthcare illustra NAP-5 Columns (Fisher 45-000-151)
5. Column equilibration and elution buffer: PBS with 1mM-EDTA.
6. Vivaspin-500 Polyethersulfone concentration spin column (Vivaproducts: VS0191, VS0121, VS0131, VS0141) with an appropriate molecular weight cutoff (3k-100k) of approximately half the protein molecular weight or lower.

### 2.3 Cell plating and sample preparation

Cells are prepared on glass substrates conducive to optical microscopy. Non-adherent cell lines can be attached to coverslips functionalized with adhesion molecules such as integrin ligands or poly-L-lysine. We use VCAM-1 coated dishes to promote attachment of CH27 B cells.

1. Cell culture dishes with #1.5 glass coverslips (Mattek P35G-1.5-14-C or similar)
2. (optional) Reagents to facilitate cellular adhesion to glass, e.g. VCAM-1/Fc chimeric proteins and anti-human Fc antibodies (recombinant human VCAM-1/Fc chimera protein, Fisher 862-VC-100; Fcγ-specific goat anti-human IgG, Jackson Immunoresearch 109-005-008) or poly-L-lysine (Sigma P4707).
3. (optional) oxygen plasma cleaner (e.g. Harrick PDC-32G)

### 2.4 Imaging reagents

1. Fiducial markers: sub-diffraction registration of multiple color channels requires reference markers that fluoresce in both channels, such as small multiwavelength fluorescent beads (e.g. .1µm Tetraspeck beads, ThermoFisher T7279). We prepare a fiducial marker sample by first cleaning a #1.5 glass coverslip cell culture dish in an oxygen plasma cleaner at full power for 3 min. A 1:100 dilution of beads in PBS is immediately added to the dish and incubated for 30 min-2 hr at room temperature. Unbound beads are removed with gentle rinsing. This sample can be stored in PBS at 4°C and reused.
2. Imaging buffer reagents: live cells are imaged in buffer conditions that maintain the cellular signaling process under study and are also conducive to single molecule photoswitching necessary for localization imaging of organic dyes. Different formulations are possible and generally include a reducing agent and oxygen scavenging system to support dye photoswitching. We use the following reagents for SiR/mEos3.2 two-color imaging:
  1. L-glutathione reduced (Sigma G6013) (see Note 1).
  2. Catalase stock solution: 4mg/ml in PBS (see Note 1).
  3. Glucose oxidase stock solution: 10mg/ml in PBS (see Note 1)
  4. Imaging buffer (30 mM Tris, 9 mg/ml glucose, 100 mM NaCl, 5 mM KCl, 1 mM KCl, 1 mM MgCl2, 1.8 mM CaCl2, 10 mM glutathione, 8 µg/ml catalase, 100 µg/ml glucose oxidase, pH 8) (See Note 1).
  5. Streptavidin or other reagents to cluster cell surface proteins or receptors, antigens to stimulate cell signaling responses, etc.

### 2.5 Microscope and imaging software

1. Many available commercial systems that are designed for single molecule imaging are compatible with super-resolution localization microscopy. In general, relatively high-power laser illumination is needed for imaging of organic dyes and multiple laser lines are needed for multicolor imaging. To image the plasma membrane selectively, illumination is set up in a total internal reflection (TIR) configuration. A high numerical aperture (NA) objective lens is used to maximize photons collected from single molecules. Multiple color channels are split using emission splitting optics and recorded a high sensitivity EMCCD or sCMOS camera. The camera control computer and software must be capable of acquisition and storage of a high volume of image data. Acquisition software should record image metadata including acquisition timestamp information. For our setup, we use an Olympus IX81 inverted microscope outfitted with the following:
  1. Coherent OBIS fiber-pigtailed solid state lasers: 405nm (50mW), 561nm (120mW), and 647nm (100mW)
  2. Olympus cellTIRF module to adjust laser incident angle
  3. 100X UAPO TIRF objective (NA = 1.49)
  4. LF405/488/561/647 quadband filter cube (TRF89902, Chroma)
  5. Olympus active Z-drift correction (ZDC) module
  6. DV2 (Photometrics) or Gemini (Hamamatsu) emission splitter.
  7. Andor iXon-897 EMCCD camera

### 2.6 Post-processing and data analysis

1. Software and computing resources are needed for post-processing of raw image data. Handling large amounts of image data and determining single-molecule localizations from raw data are computationally intensive. Numerous commercial and freely available software packages are available for single molecule localization. We find that the imageJ plugin ThunderSTORM [38] is flexible and accessible. We use MATLAB for all further analysis of single molecule localization to quantify protein co-localization and dynamics.

## 3. Methods

All steps are carried out at room temperature unless otherwise noted.

### 3.1 Preparation of fluorescent Fab fragments

Labeling proteins such as antibodies, Fab fragments, toxins, or other binding proteins can be conjugated with organic dyes suitable for super-resolution imaging (see Note 2) via NHS ester chemistry. In these reactions, a reactive group commonly available on organic dyes is used to covalently attach the dye to amines on lysine residues or N-termini of purified proteins. After conjugation, unreacted dye is separated from labeled protein by size exclusion. For applications where the labeling protein is cross-linked during the imaging experiment, for example to study the effects of protein clustering, labeling proteins can be conjugated with both dye and biotin groups using the same succinimidyl ester chemistry. In our specific application, we label goat anti-Mouse IgM(µ) Fab fragments with biotin and silicon rhodamine (SiR) NHS esters.

1. Conjugation reactions should be carried out near pH 8.5 under buffer conditions where proteins are stable. Exchange purified proteins into the reaction buffer through dialysis. If the initial protein storage buffer contains free amines such as Tris or glycine, several rounds of dialysis may be necessary to remove them. Final protein concentrations between .5 – 2mg/ml are optimal for labeling.
2. Add amine-reactive dye succinimidyl ester DMSO stock solutions to labeling protein solutions in 2 - 8x molar excess (see Note 3). Immediately mix the protein solution by inversion or by flicking the tube to avoid exposing proteins to high local concentrations of DMSO.
3. Incubate the reaction for 1 hr at room temperature with rotation. Protect from light e.g. by enclosing the reaction tube in a larger foil-covered tube.
4. After incubation, pass the reaction solution through a NAP-5 gel filtration column using PBS with 1mM-EDTA as the equilibration and elution buffer.
5. Modified proteins can be further purified and concentrated by centrifugation in a Vivaspin-500 Polyethersulfone concentration spin column.
6. Estimate the degree of labeling using absorbance measurements of the dye-conjugated protein at the dye’s absorption maximum and 280nm. We aim for a dye-to-protein ratio around 2. If the protein is under-modified, the reaction can be repeated to increase the dye-to-protein ratio.
7. For proteins modified with both fluorescent dyes and biotin, we use a two-step procedure. First, react unlabeled proteins with a mixture of fluorophore and biotin NHS esters in 5x and 12x molar excess, respectively. Purify as described above. Second, repeat the labeling procedure with only fluorophore NHS ester in 5x molar excess.

### 3.2 Cell preparation

Cells can express tagged proteins e.g. through transient transfection or through the preparation of stable cell lines through antibiotic selection or viral transduction. Expressible fluorescent tags can take the form of photo-activatable/photo-switchable fluorescent proteins or self-labeling protein tags that covalently bind to organic dyes such as SNAP tags and Halotags. Proteins on the cell surface can be labeled with antibodies or toxins conjugated to organic fluorophores (see 3.1). Labeling strategies that use transient binding and exchange of fluorescent conjugates to achieve stochastic blinking, as in PAINT and DNA-PAINT, are also compatible. Our most common two-color labeling approach is to pair BCR labeling using SiR-biotin-conjugated Fab fragments with transient expression of an mEos3.2 fusion protein. Prior to imaging, cells should be plated on dishes or wells with #1.5 glass coverslips under conditions that facilitate uniform adhesion.

1. If needed, transiently transfect cells with fluorescent fusion constructs and allow them to recover in growth media in the 37°C CO_2_ incubator for 18-24 hours before preparation for imaging.
2. If needed, treat glass bottom wells or culture dishes with reagents to facilitate adhesion. For example, VCAM-1 coated dishes can be prepared as follows:
  1. Plasma clean imaging chambers using an oxygen plasma cleaner at the highest power setting for 3 min.
  2. Immediately incubate cleaned dishes with a solution of Fcγ-specific goat anti-human IgG antibodies in PBS at 100 μg/mL for 30 min at room temperature.
  3. Rinse dishes with PBS and block for 30 min with a 5% BSA solution in PBS.
  4. Rinse dishes with PBS and incubate dishes for 1 hour at room temperature or overnight at 4°C with recombinant VCAM-1/human Fc chimera protein at 10 μg/mL,
  5. Rinse dishes thoroughly in PBS before use.
  6. Coated dishes can be stored in PBS at 4°C for up to 1 week.
3. Cells can be grown overnight on untreated glass, but should be plated soon before imaging if 1) cells secrete an expressed fluorescent protein which can become deposited on the glass (see Note 4) or 2) a surface treatment such as VCAM-1 coating is used. CH27 cells adhere to untreated glass within 2-4 hours at 37°C and VCAM-1 coated dishes within 10 min at room temperature.
4. In both cases, cells should be plated at a density such that they remain isolated from one another for imaging.
5. A day of imaging usually involves multiple replicates of a live cell imaging measurement. Prepare as many dishes of cells as replicates are desired.
6. Label cells with dye conjugated proteins at room temperature or on ice if needed to minimize probe internalization. We label BCR by incubation with 5 µg/ml SiR-biotin-Fab in culture medium for 10 min at room temperature. Cells should be labeled shortly (within 30 min) before imaging to prevent internalization.
7. At this stage and prior to imaging, cells can be pre-treated with drugs or other perturbations as needed for the experiment.

### 3.3 Imaging

Sparse, single molecules are simultaneously imaged in two color channels over time.

1. Configure microscope such that two fluorophores can be imaged simultaneously. This likely will involve illumination from several excitation lasers (e.g 405nm, 561nm, 647nm) paired with a multi-band filter cube that reflects laser lines and passes fluorophore emission. An emission splitter unit is used to separate emission from distinct fluorophores and direct them onto separate cameras or separate sides of the same EMCCD or sCMOS camera (see Note 5). Lasers and emission optics should be aligned prior to sample preparation. We recommend using an automated correction for drift in the axial direction (e.g. ZDC from Olympus). Total internal reflection (TIR) illumination is commonly used for selective imaging of cell surface proteins and structures on the plasma membrane.
2. Acquire images of fiducial markers to register color channels: Take many (>100) images of the fiducial marker bead sample using the same illumination laser lines and filters to be used in the cellular imaging measurement. Between acquisition of individual bead images, translate the stage so that the field of view is uniformly sampled by bead positions. It is important that the splitting optics are stable throughout the course of the imaging experiment. We recommend acquiring bead images before and after a cell measurement, to ensure that the relative positions of the two color channels do not shift during a measurement, impairing channel registration.
3. Add oxygen-scavenging enzymes glucose oxidase and catalase to the imaging buffer (see Note 1) and replace buffer in cells with fresh imaging buffer.
4. Select cells for imaging. Cells should be firmly adhered to the dish to minimize membrane topography. It can be helpful to image cells using an interference reflection microscopy (IRM) [39] filter cube to screen for uniform membrane flatness across the cell footprint. Alternately, dSTORM probes can be viewed with low laser power with TIR illumination to assess the uniformity of the signal **(Figure 1A)**, or the blue excitation/green emission of un-converted mEos3.2 can be used for the same purpose. Cells should also be chosen based on expression levels of fluorescent constructs or labeling density.
5. Adjust laser intensities and frame rate. The goal for imaging is to view sparse single molecules in each color channel **(Figure 1B)** over an extended period of time (10min). High laser intensity is needed to induce dSTORM probe photoswitching and the probe density is inversely dependent on illumination intensity. 405nm laser intensity can be used to control mEos3.2 probe density (See Note 7). For SiR/mEos3.2 imaging, we use the following range of laser powers during single molecule data collection: 5-20 kW/cm^2^ for 561 nm and 647 nm lasers, and 100-200 W/cm^2^ for the 405 nm laser. Frame-rates should be chosen such that exposure times are as long as possible to maximize signal-to-noise but short enough to minimize motion blur and accurately localize mobile single molecules. Exposure times in the tens of milliseconds corresponding to frame rates between 20 and 100 Hz are typical, depending on the capabilities of the camera and mobility of proteins of interest.
6. Acquire single molecule images over time. Acquire several minutes of images of the untreated cell prior to carefully adding a treatment or activator (see Note 6). We add streptavidin during the experiment to cluster SiR-biotin-Fab labeled BCR and induce cellular activation. Image acquisition timestamps can be useful for keeping track of changing behavior relative to the time of treatment addition. When adding treatments during imaging, add relatively large volumes to facilitate fast mixing. Add treatments diluted in imaging buffer. We recommend saving images in successive multistack or movie files with a set number of images per file (e.g. 500 frames corresponding to 10s). Smaller file size speeds saving during acquisition and loading during post-processing.
7. Independent measurements should be repeated over multiple days using multiple biological replicates.

### 3.4 Post-processing: single molecule localization

Single molecules can be localized using a wide variety of software packages. The end goal is to generate a list containing the positions and other attributes of single molecules over time in each color channel.

1. Localize single molecule positions. This can be accomplished in a wide range of software packages, including imageJ using the Thunderstorm plug-in [38]. This particular software package has implemented multi-emitter fitting, which allows for the accurate processing of images with a high density of single molecules per frame. Localized positions and other fit parameters can be exported for further processing (See Notes 7 and 8).
2. Generate a spatial mapping between image channels using fiducial markers, for example using the procedures described in [40]. To accomplish this, fiducial markers are first matched across image channels and used as control points to generate a transform. Once the positions of the control points are established, this map can be generated from control points using the function fitgeotrans() and it can be applied to points using the function transformPointsForward() in MATLAB.
3. Cull localizations to remove poor fits. This is typically done by accumulating distributions of fit parameter values and removing the top and bottom 5% for parameters such as intensity or width.
4. Localized positions can be corrected for stage drift or collective cellular motions using a drift correction algorithm, for example using the procedures described in [41]. Although drift correction is not strictly required for the cross-correlation or diffusion analysis because it applies at slower time-scales than are typically of interest, it can be important when establishing a region of interest (ROI) that is appropriate for localizations acquired over the entire dataset.
5. Images representing the time-averaged positions of localized molecules can be rendered by generating two dimensional (2D) histograms from the localized positions. This can be accomplished using the histcounts2 function in MATLAB. Histograms can be presented as is or blurred using a Gaussian filter for display purposes.

### 3.5 Post-processing: Cross-correlation analysis

Cross-correlation analysis is conducted using the locations of molecules in both space and time. Cross-correlation functions are generated by tabulating pairwise distances between localizations in different color channels, assembling these distance into histograms, then normalizing these histograms to account for the expected number of pairs if probes are randomly co-distributed over the specified region of interest. We have assembled a collection of MATLAB functions to perform these calculations from lists of localizations, as well as sample analysis scripts, publically available at https://github.com/VeatchLab/smlm-analysis. Scripts and localizations used to generate **Figures 2-5** are also included in the SMLM-analysis distribution.

These methods derive their power from the assumption that the effective interactions between labeled proteins are similar at different locations across the region of interest, so that localizations from different locations on the cell can be thought of as samples from one statistical distribution. By combining all of these observations, the poor sampling at any one location adds up to excellent sampling of the pair distribution overall. Note that this assumption may not hold for some structures of interest – for example a polarized cell may give rise to different effective interactions on the two poles.

**Figure 1:**
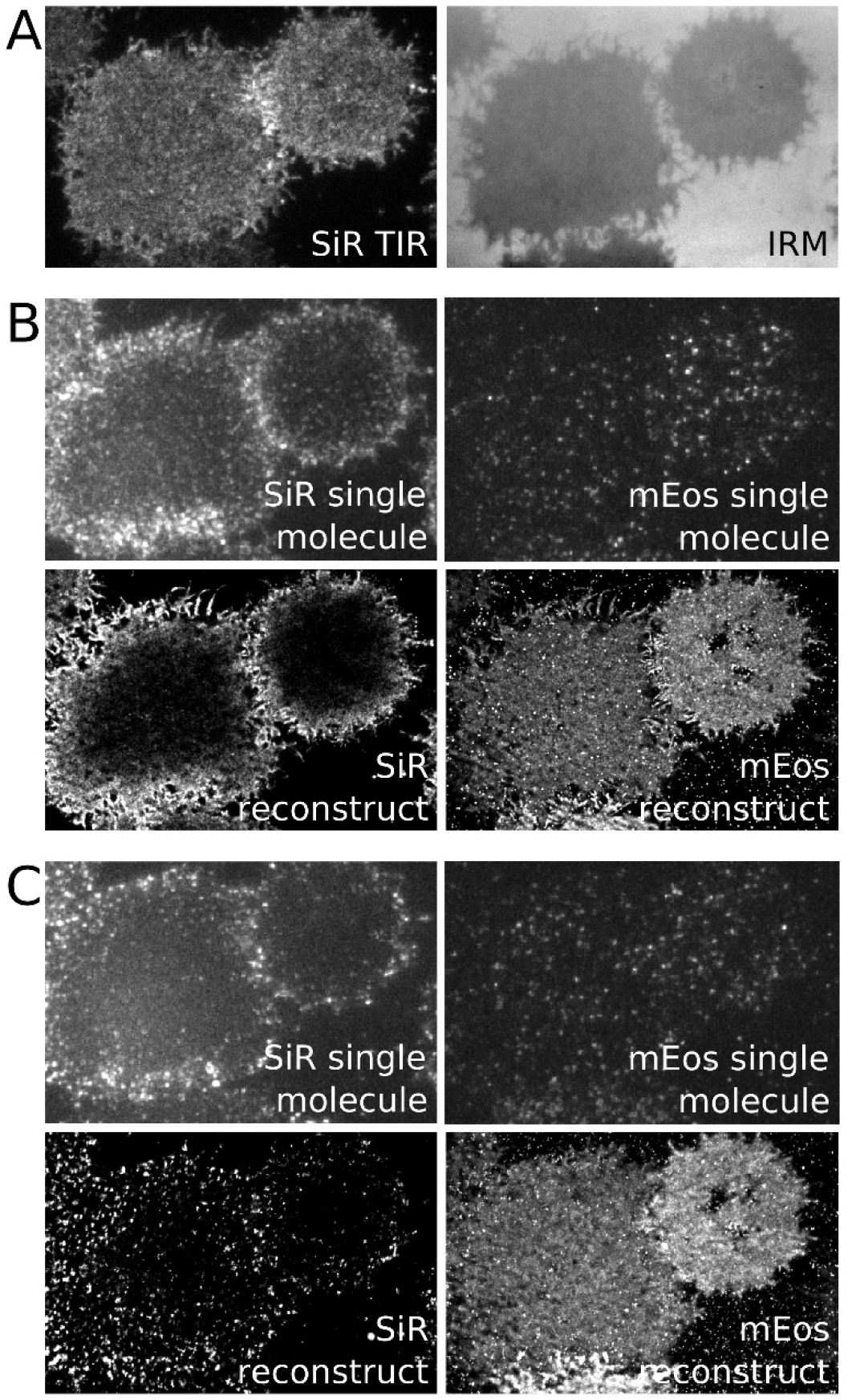
Sample raw and reconstructed images. (A). Total internal reflection (TIR) and interference reflection microscopy (IRM) images of an adherent CH27 B cell after incubation with a Fab fragment αIgMμ conjugated to both SiR and biotin. When imaged with low intensity 647nm excitation, SiR should exhibit a uniform and bright distribution over the cell surface when imaged in TIR and the IRM image should also be uniform indicating minimal membrane topography. (B) (top) Raw TIR image frames of SiR imaged at high 647nm laser power and mEos3.2 imaged with both 561nm excitation and 405nm activation lasers. Imaging SiR with high laser power converts some probes into a reversible dark state allowing the visualization of isolated single molecules. (bottom) Super-resolution image of SiR and mEos3.2 from localizations acquired from 4500 image frames over 2.9 min. Note the presence of a gradient in the SiR image. This is due to mobile probes diffusing from the cell edges that are not initially in a reversible dark state since the TIR field does not extend above the ventral cell surface. (C) The same conditions as B but images are acquired after the addition of streptavidin to induce clustering and activation of BCR. Super-resolved images are generated from 8500 raw images frames acquired over 5.8 min.

**Figure 2:**
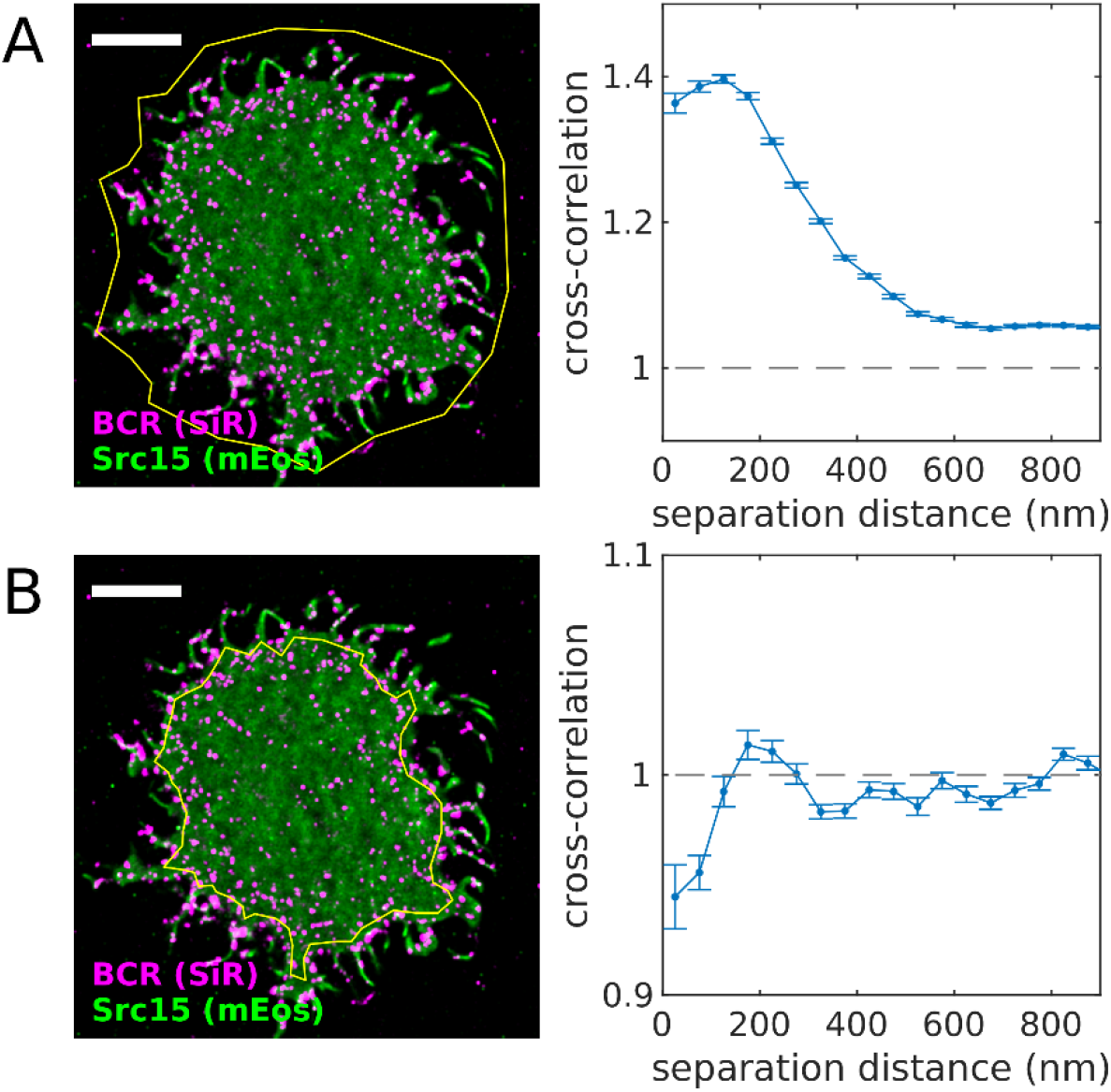
Defining an appropriate region of interest (ROI). (A) Reconstructed image (left) of SiR-tagged BCR and transiently-expressed Src15-mEos3.2 in an adhered CH27 B cell imaged after BCR clustering with streptavidin. The yellow polygon indicates the region of interest (ROI) used for the cross-correlation analysis (right). This ROI contains areas inside and outside the cell footprint, including membrane topographical features of the cell boundary itself and thin protrusions. Because both probes reside on the cell membrane, the cross-correlation takes on values > 1, indicating colocalization of the two probes. (B) (left) The same cell shown in (A) with a ROI defined within the cell boundary. (right) The cross-correlation function tabulated using this ROI indicates exclusion of the two probes (values < 1), reflecting the co-distribution of these proteins within the membrane. A MATLAB script that generates this figure is included in the SMLM-analysis distribution.

Functions within this publically available package are referenced below.

1. Specify the region of interest (ROI) for further analysis. ROIs should be placed within the cellular footprint and not contain cell edges. ROIs should also exclude other clear topographical features. Since localizations are analyzed as projections on two dimensions, features that contain more membrane per area (e.g. cell edges, filopodia) will lead to trivial correlations. A tool to draw ROIs on reconstructed images called spacewin_gui() is included in the SMLM-analysis distribution. This tool allows a user to draw polygons on images reconstructed from lists of localized positions. Polygons can be drawn that either include or exclude points, allowing for ROIs with complicated topologies. Examples of different ROIs and their impact on the resulting cross-correlation functions are shown in **Figure 2**.
2. Tabulate cross-correlations that properly normalize for ROI boundaries. This can be accomplished using the MATLAB function spacetime_xcor() posted in the SMLM-analysis distribution. This base function includes an efficient code to tabulate the pairwise distances between localizations over time, adapted from [42] and described previously [41], as well as boundary conditions for arbitrary spatial ROIs and temporal windows, as described [34]. In some cases, long-range gradients in images can obscure results when an analysis of short-range correlations are desired. To reduce the impact of long-range gradients, a local density correction can be applied, tabulated using spatial_gradient_correction() in the SMLM-analysis distribution, following an approach described previously [43]. We find that this correction is often necessary when imaging mobile (d)STORM probes such as SiR that diffuse into the illuminated area from the cell edges in an activated state. These probes tend to take some time to be converted into a reversible dark state, leading to a larger density of probes at the cell periphery (evident in **Figure 1B** for SiR). A properly normalized correlation function should decay to 1 when probed at large separation distance or large separation-times. Examples of correlation functions tabulated from the same set of localizations but with different normalizations are shown in **Figure 3**.
3. Correct for bleed-through and/or other fluorophore related photo-physical effects between image channels. Fluorophores have broad emission spectral characteristics, making it difficult to fully isolate emission with filters alone. Even low intensity emission in the incorrect color channel can bias the localizations to generate the appearance of shorter pairwise distances, when molecules appear in both color channels simultaneously and within a diffraction limited separation distance [31]. FRET or other proximity related photo-physical effects (e.g. [44]) can also give rise to correlations that do not reflect the actual density of labeled molecules. These effects can be detected and corrected by analyzing how cross-correlations vary with the time-interval between observations. Bleed-through or other fluorophore related photo-physical effects are expected to be most apparent when probes are observed simultaneously within a diffraction limited separation distance (∼200nm). An example is shown in **Figure 4** in which one probe (SiR labeling BCR) is clustered and largely immobile and the second probe (mEos3.2 labeling a minimal GPI linked protein) is mobile. In this case, cross-correlation amplitudes (**Figure 4B**) are highest for simultaneous observations (time-separation = 0) and decay over a fraction of a second for short separation distances (r<150nm). Since BCR clusters are largely immobile, the cross-correlation amplitude should not depend on when exactly labels decorating those clusters are observed, therefore we expect the actual cross-correlation amplitude to not vary with time-interval. These artificial probe-related correlations can be removed by either averaging over a range of time-intervals excluding the effected points, or by fitting quickly varying contributions and reporting only the slowly varying component. The impact of these corrections can be seen in the spatial correlation functions of **Figure 4C**.
4. In some cases, we are most interested in the change in the magnitude of correlations upon stimulation, in which case we subtract correlation functions generated from localizations detected before stimulation from the correlation function generated from localizations obtained after stimulation (see Note 6). One advantage of this approach is that it provides a means to exclude systematic contributions from membrane topography that are present over the entire measurement but are not removed by the ROI. We found this to be important when quantifying small signals [45].

### 3.6 Post-processing: Diffusion analysis

Single molecule mobility can be characterized by quantifying the evolution of the spatial auto-correlation in time. As molecules move, they tend to explore larger distances over larger time-intervals, and this can be detected as a broadening of the initial peak in the auto-correlation function, over time-scales relevant to the off-rate of the fluorophore blinking. This analysis yields similar information as is obtained in single particle tracking, but has the advantage of not requiring that localizations be assigned to trajectories, which may be ambiguous for densely sampled data. An example showing GPI-mEos3.2 motion is shown in **Figure 5**. The script and localizations used to generate Figure 5 is included in the SMLM-analysis distribution.

**Figure 3:**
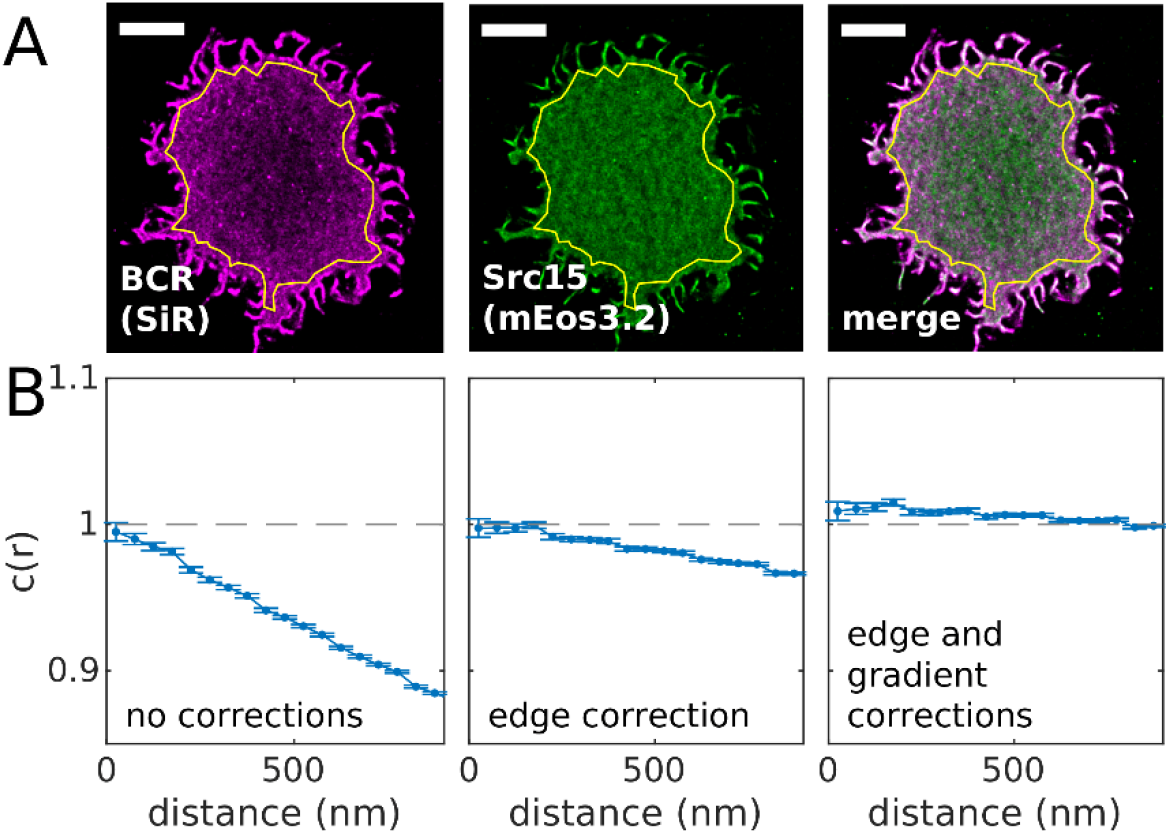
The normalization of the cross-correlation function c(r). (A) Reconstructed image of SiR tagged BCR and Src15-mEos3.2 in an adherent CH27 B cell imaged before BCR clustering with streptavidin. The ROI used is shown in yellow. (B) Cross-correlation functions tabulated for localizations within the region of interest in A. The three curves shown contain different corrections to the cross-correlation normalization as indicated. The edge correction incorporates the geometry of the region of interest to account for the reduced number of possible pairs for localized molecules close to edge of the ROI. The gradient correction accounts for changes in local density that vary over length-scales greater than 1μm in this example. A MATLAB script that generates this figure is included in the SMLM-analysis distribution.

**Figure 4:**
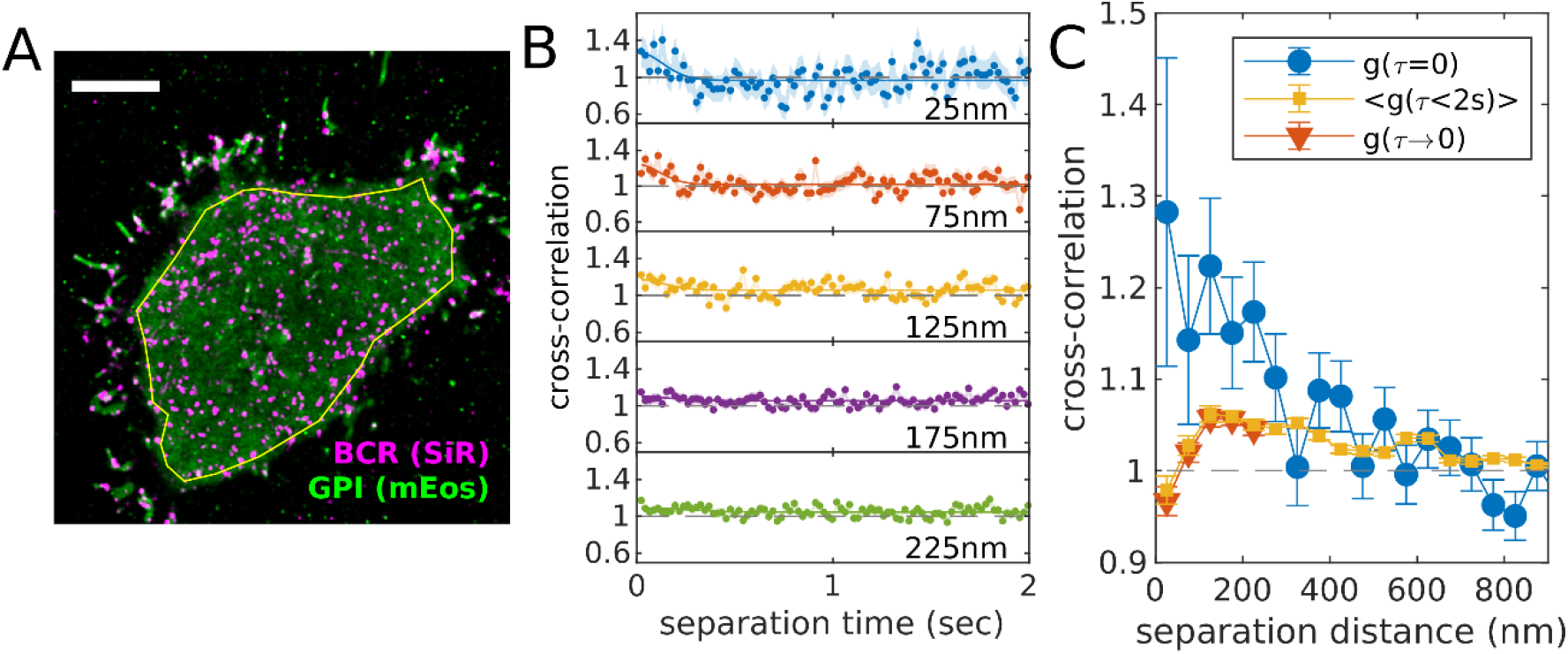
Example of procedure to account for fluorophore bleed-through or other temporally correlated photo-physical effects. (A) Reconstructed image of SiR tagged BCR and GPI-mEos3.2 in an adherent CH27 B cell imaged after BCR clustering with streptavidin. The yellow polygon indicates the ROI used for the cross-correlation analysis. (B) Cross-correlation functions plotted as a function of separation time for several spatial bins centered at the values indicated. BCR clusters are largely immobile over these time-scales (2 sec), therefore a uniform or slowly-varying cross-correlation amplitude is expected in the absence of bleed-through or other photo-physical effects. We attribute the upwards inflection of cross-correlation curves at short separation times to fluorophore-dependent effects since they occur over diffraction-limited spatial scales (<200nm) and diffusion-limited time-scales (<0.25sec in this example). (C) This artifact results in the appearance of correlations that are apparent if only simultaneously-observed localizations are included (g(τ=0)). This artificial correlation can be removed by either reporting the average over all time-intervals (<g(τ<2sec)>) or by fitting the time-interval dependent upwards deflection in g(τ) to a Gaussian shape (solid lines in B) and removing this component from the residual function extrapolated to τ=0 (g(τ→0)). A MATLAB script that generates this figure is included in the SMLM-analysis distribution.

**Figure 5:**
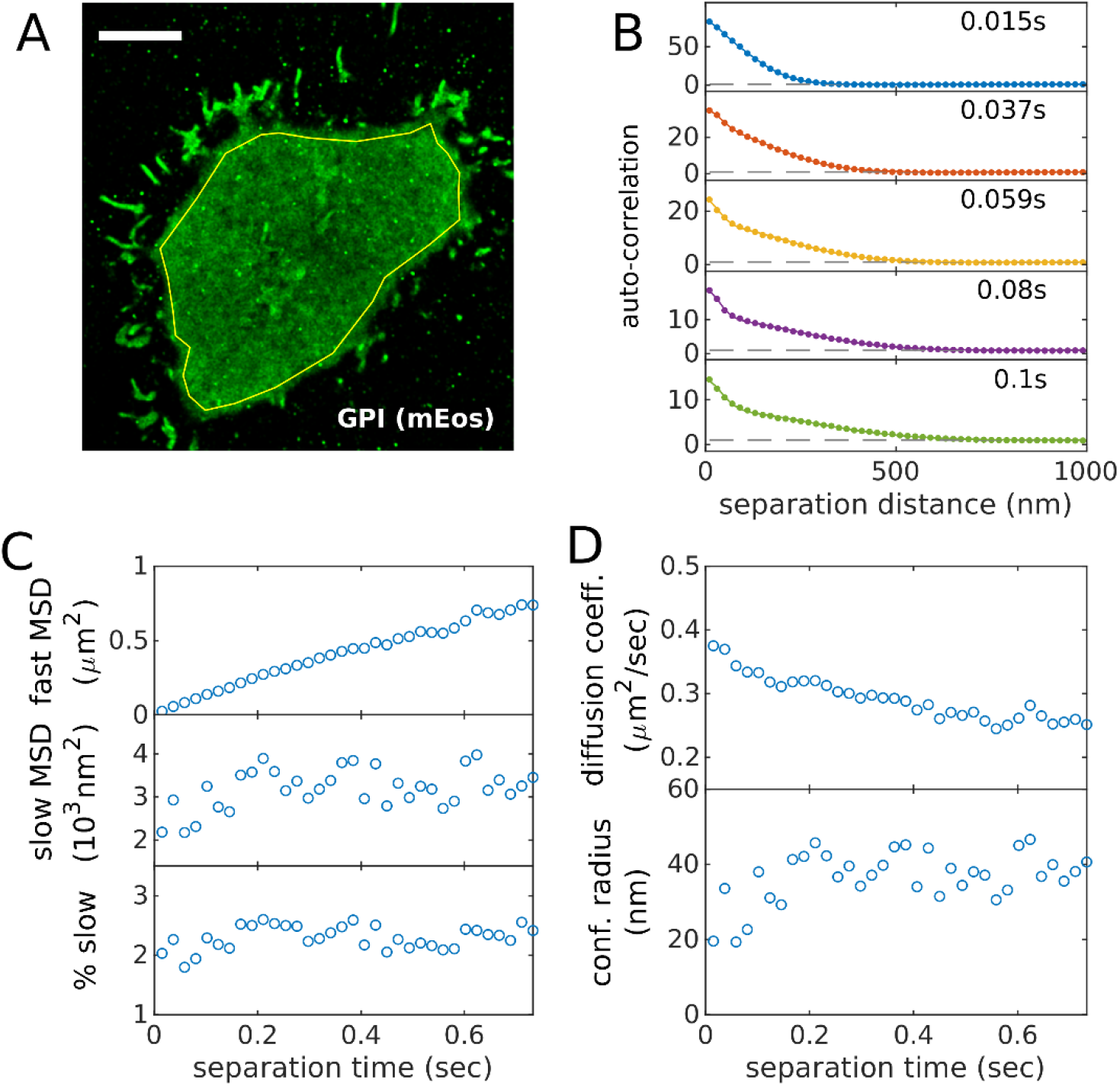
The time-dependent auto-correlation function quantifies protein mobility. (A) Reconstructed image of GPI-mEos3.2 in an adherent CH27 B cell imaged after BCR clustering with streptavidin. The yellow polygon indicates the ROI used for the auto-correlation analysis. (B) Auto-correlation functions plotted as a function of separation distance for several temporal bins centered at the values indicated. Curves are fit to a superposition of two Gaussian functions to capture the two populations. As the time-interval is increased, the faster population extends to larger separation distances while the slower population remains confined to short separation distances. (C) Parameters extracted from fits like those shown in B including the mean squared displacement (MSD) for the fast and slow populations as well as the % of segments associated with the slow state. In this case, the fast MSD is roughly linear in time-interval while the slow MSD and the fraction of segments in the slow state does not change meaningfully with time-interval. (D) Mobility parameters evaluated from fit parameters in C. A MATLAB script that generates this figure is included in the SMLM-analysis distribution.

1. Using the same ROI constructed for the cross-correlation analysis, tabulate the spatial auto-correlation for the color channel corresponding to the molecule being characterized. This can be done using the same MATLAB function used to tabulate cross-correlations, spacetime_acor(), which resembles the function spacetime_xcor() but in this case only positions for one color channel are inputted. Auto-correlation functions should include normalizations for the ROI and spatial gradients.
2. Auto-correlation functions can be fit to a superposition of Gaussian functions to extract the mean square displacements (MSDs) and relative populations associated with each state. The example of Figure 5 is fit to a superposition of two Gaussians of the form:

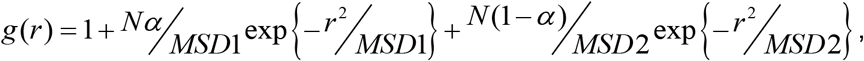

where *N* is a normalization factor, *MSD1* and *MSD2* are the mean squared displacements in the two states, and *α* is the fraction of molecules in the 1^st^ state. This fitting approach makes the assumption that motion is normally distributed, which is typically appropriate as long as directed motion is not expected. This approach does not assume that motion is Brownian.
3. Best fit MSDs can be converted to diffusion coefficients at a given time-interval τ according to *D* = *MSD* 4*τ*. Note it is important to account for the finite integration time and finite localization precision of the measurement, especially for short time-intervals [46]. The effective time-interval is calculated as 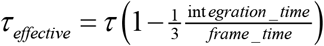 and the effective mean squared displacement is *MSD*_*effective*_ = *MSD* − 2(*LP*)^2^ where *LP* is the average localization precision of the measurement. For the example of Figure 5, the slow state is highly confined, in that its MSD does not increase with time-interval. In this case it is more appropriate to report the confinement radius, which is 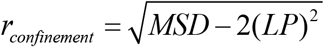

## 4. Notes

1. Some steps should be taken to ensure the maximal activity of the imaging buffer. Glutathione powder should be stored at 4°C under desiccation and allowed to come to room temperature before weighing to avoid oxidation. Glutathione is unstable in aqueous solutions so imaging buffers should be prepared on the day of the experiment. Catalase stock solutions have a poor shelf life and should not be frozen, and are therefore best prepared within 48 hours of use and stored at 4°C. We typically prepare imaging buffer stock solutions containing all components except for glucose, glutathione, glucose oxidase, and catalase for long-term storage at 4°C. On the day of the experiment, glucose and glutathione are added and the pH of the buffer is adjusted to 8 with small amounts of 10M NaOH to counteract the acidifying effect of glutathione. Glucose oxidase and catalase are added immediately before imaging. Final concentrations of glucose oxidase and catalase strongly effect the blinking kinetics of organic dyes and need to be optimized for the specific dyes being used as well as parameters such as labeling density. The concentrations given are a useful starting point.
2. For quantitative measurements of protein co-distributions by super-resolution imaging in two color channels, special considerations apply for dye selection. It is imperative that each orthogonal label is identified in one color channel only. Cyanine-based dyes commonly used in localization microscopy can undergo reactions that shift their absorbance spectra to shorter wavelengths, leading to detection of spurious signals in the wrong color channel from these dyes [47, 48]. Two-color detection in the red and far-red emission channels is commonly used because of availability of dyes and fluorescent proteins optimal for localization microscopy. For this combination, we favor silicon-substituted rhodamine dyes [49] for far-red detection.
3. Molar ratios of succinimidyl ester dye to target protein used for dye conjugation vary based on the specific dyes and proteins and need to be optimized empirically. Over-modification of proteins with organic dyes can lead to protein aggregation and instability in solution. In our experience it is preferable to begin with lower molar ratios and perform multiple successive conjugation reactions if necessary.
4. We have found that some transiently expressed fluorescent fusion proteins are secreted by cells. These proteins can become deposited on glass coverslips, resulting in immobile mEos3.2 molecules that are difficult to distinguish from tagged proteins of interest in the intact plasma membrane. Deposited proteins accumulate over time and can be minimized by plating cells on coverslips for a few hours after they have recovered from transfection.
5. Quantitative analysis of multi-color SMLM data is very sensitive to even low levels of spectral bleed-through across color channels. Emission filters should be selected to minimize bleed-through, with more restrictive requirements than are typical for standard TIR or wide field fluorescence imaging. Small residual bleed-through artifacts can be corrected in post-processing (see 3.5.3 and Figure 4).
6. Factors such as subtle membrane topography can contribute to cross-correlation functions in ways that can be difficult to identify or isolate (see 3.5.1 and Figure 2). These contributions add to cell-to-cell variation in cross-correlation functions and are not typically the focus of the imaging experiment. In situations where structures are expected to be present throughout the imaging experiment, it can be useful to quantify the difference in the cross-correlation function before and after the addition of a cell treatment or stimulus rather than the absolute amplitude of the cross-correlation after treatment. For this reason, it can be helpful to acquire sufficient image data before treatment to establish a baseline for the cross-correlation function.
7. Quality assessment for raw data requires special attention. It should be noted that fitting algorithms are capable of producing localizations from even very poor quality raw data and therefore the ability of processing procedures to reconstruct super-resolution images is not a sufficient data quality standard. Isolated single molecules should be identifiable by eye in raw images. Reconstructed images generated from raw data with a high density of single molecules often contain recognizable artifacts such as intensity bleeding between higher intensity objects or narrowing of linear structures. Localization algorithms return quality control metrics (e.g. localization error, spot width and intensity, etc.) that degrade when probe density is too high, indicating single molecules are not well resolved.
8. It can be useful to localize single molecules on images that have undergone a background subtraction to remove intensity from signals that do not vary in time. The background is typically estimated by taking the median of images acquired over some finite time (e.g. 500 frames), subtracting this from the raw images, then adding back the average intensity value of the median image. The most commonly used localization algorithm [50] incorporates information about pixel intensity, camera gain and noise when localizing and estimating the localization error on single fits. These error estimates will contain systematic errors when applied to background subtracted image frames.

## Notes

https://github.com/VeatchLab/smlm-analysis

### Competing Interest Statement

The authors have declared no competing interest.

